# Double Domain Swapping in Human *γ*C and *γ*D Crystallin Drives Early Stages of Aggregation

**DOI:** 10.1101/2020.07.18.210443

**Authors:** Balaka Mondal, Jayashree Nagesh, Govardhan Reddy

## Abstract

Human *γ*D (H*γ*D) and *γ*C (H*γ*C) are double domained crystallin (Crys) proteins expressed in the nucleus of the eye lens. Structural perturbations in the protein often trigger aggregation, which eventually leads to cataract. To decipher the underlying molecular mechanism, it is important to characterize the partially unfolded conformations of Crys proteins. Using coarse grained protein models and molecular dynamics simulations, we studied the role of on-pathway folding intermediates in the early stages of aggregation. The multi-dimensional free energy surface revealed at least three different folding pathways with the population of partially structured intermediates. The two dominant pathways confirm sequential folding of the N-terminal [Ntd] and the C-terminal domains [Ctd], while the third, least favored pathway involves intermediates where both the domains are partially folded. A native like intermediate (*I*^∗^), featuring the folded domains and disrupted inter domain contacts, gets populated in all the three pathways. *I*^∗^ forms domain swapped dimers by swapping the entire Ntds and Ctds with other monomers. Population of such oligomers can explain the increased resistance to unfolding resulting in hysteresis observed in the folding experiments of H*γ*D Crys. An ensemble of double domain swapped dimers are also formed during refolding, where intermediates consisting of partially folded Ntds and Ctds swap secondary structures with other monomers. The double domain swapping model presented in our study provides structural insights into the early events of aggregation in Crys proteins and identifies the key secondary structural swapping elements, where introducing mutations will aid in regulating the overall aggregation propensity.

Graphical Abstract figure

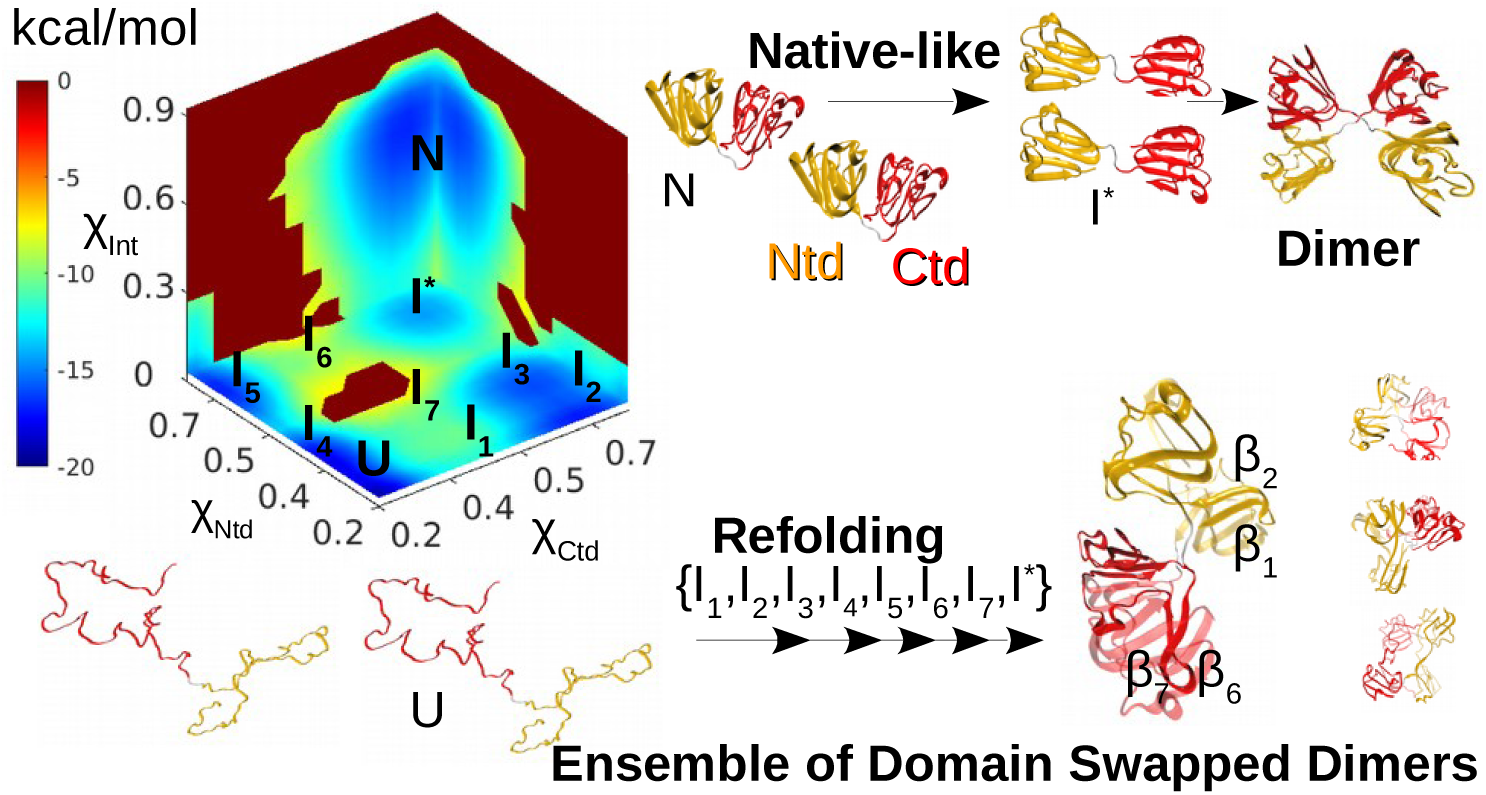

## Introduction

Lens opacity or cataract[1] is a protein aggregation disorder associated with misfolding and subsequent aggregation of eye lens crystallin (Crys) proteins, resulting in partial or complete loss of vision. Detailed structural characterization of the misfolded states and molecular mechanism of cataract formation have remained elusive to date. In humans, surgically removed cataractous lenses contain high molecular weight aggregates of *α, β* and *γ* Crys, the three main classes of Crys found in vertebrates[2, 3]. *α* Crys primarily acts as molecular chaperones[4], while *β* and *γ* Crys are responsible for optical functions of the lens[5, 6].

Monomeric *γ* Crys are the simplest members of the Crys family. The three most abundant *γ* Crys proteins in human are H*γ*C, H*γ*D and H*γ*S Crys, which are present in the central region of the lens. H*γ*C and H*γ*D Crys are expressed in the lens nucleus[7], which is the oldest region of the lens with no protein turnover. Since these proteins are only synthesized once, the native state of these proteins should be stable and functional for human lifetime. Any misfolding significantly perturbs the equilibrium concentrations of H*γ*C and H*γ*D Crys in the lens.

H*γ*C and H*γ*D Crys are both 173 residue long proteins with ≈ 71 % sequence similarity. The native structure of *γ* Crys is topologically complex with a N-terminal domain (Ntd) and a C-terminal domain (Ctd), which are highly homologous. Each domain consists of two intercalated greek-key motifs, which are complex supersecondary structures formed by four antiparallel *β* strands (Figure 1A,2A). Intercalated greek key motifs termed *M*_1_ and *M*_2_ constitute the Ntd, while *M*_3_ and *M*_4_ constitute the Ctd [8] (Figure 1B). A short peptide linker connects the two domains, which form a highly conserved hydrophobic interface necessary for the long term stability of the proteins [9]. The topology of the native state of *γ* Crys leads to complex folding behavior as observed in experiments[10, 11, 12, 9] and computer simulations[13, 14, 15].

**Figure 1:**
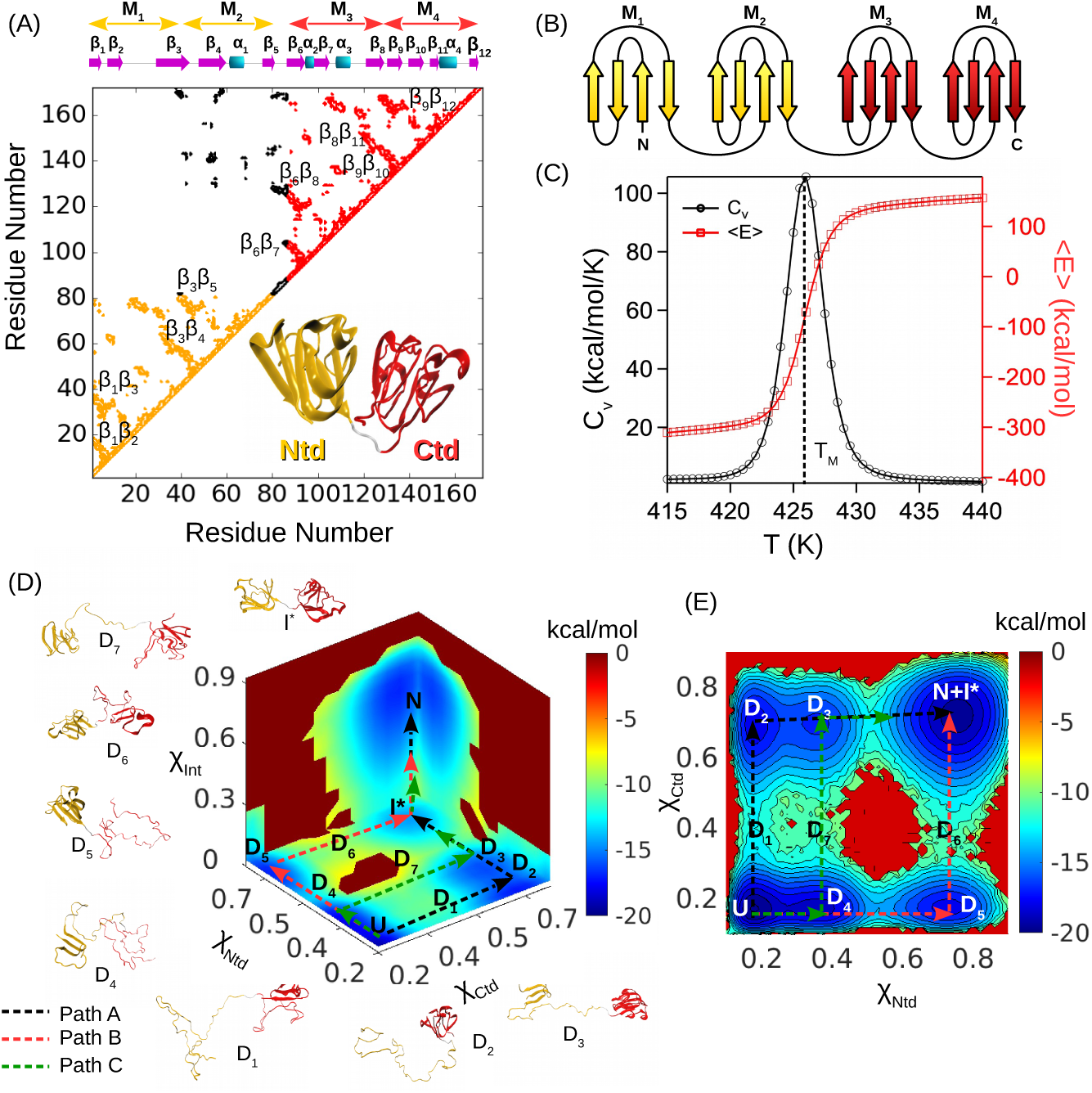
(A) Contact map of H*γ*D Crys[46] (PDB ID: 1HK0). H*γ*D Crys consists of two domains called N-terminal domain (Ntd, shown in orange) and C-terminal domain (Ctd, shown in red) connected by a short peptide linker. Each domain is further composed of two intercalated greek key motifs. Greek key motifs constituting the Ntd are termed as *M*_1_ (residues Gly1 to Ser39) and *M*_2_ (residues Gly40 to Pro82), while motifs constituting the Ctd are termed as *M*_3_ (residues His88 to Glu128) and *M*_4_ (residues Gly129 to Ile171). The intra domain contacts are colored orange (Ntd) and red (Ctd), while inter domain contacts are colored black. (B) A splay diagram of four ideal interconnected greek key motifs. Four antiparallel *β*-strands constitute one greek key motif where strands *β*_1_ and *β*_3_ are in contact. (C) Temperature dependent variation in specific heat capacity (*Cv*) and average potential energy (*(E)*) indicate a two state folding transition in H*γ*D Crys. (D) FES, *F* (*χ*_*Ntd*_, *χ*_*Ctd*_, *χ*_*Int*_) projected onto fraction of native contacts in Ntd (*χ*_*Ntd*_), in Ctd (*χ*_*Ctd*_) and between the two domains (*χ*_*Int*_) at *T*_*M*_ = 426 K. Three pathways connect the native state (*N*) and the unfolded state (*U*) through distinct intermediate states. Representative structures of the intermediates are shown. (E) *F* (*χ*_*Ntd*_, *χ*_*Ctd*_) projected onto *χ*_*Ntd*_ and *χ*_*Ctd*_. All the intermediate states are visible except for *I*^∗^ (Ctd and Ntd domains are folded with disrupted inter domain contacts), which is indistinguishable from the *N* state.

Due to their implications in cataract, biophysical properties of *γ* Crys have been studied extensively using both experimental and computational techniques. Studying folding in multi-domain proteins is challenging since the domain-domain interactions add to the complexity as they can influence folding in addition to the intra-domain interactions[16, 17, 18, 19]. Denaturant induced unfolding experiments[10, 20, 21, 11] demonstrated a three state unfolding transition in *γ* Crys indicating the presence of a partially structured intermediate state. This intermediate state is structurally characterized for H*γ*D Crys, which consists of a folded Ctd and an unfolded Ntd[22, 9], in agreement with the simulations[13].

Hysteresis is another interesting property observed in the denaturant induced equilibrium unfolding/refolding experiments[10, 23, 24]. Native states of H*γ*D Crys are observed to be stable at higher denaturant concentrations where unfolded states did not refold. Increased temperature or equilibration time led to diminished hysteresis causing a shift in the unfolding transition only, indicating a high kinetic barrier to unfolding as a possible reason for hysteresis. Further, increased stability of the inter domain interface led to enhanced hysteresis, which is absent in the truncated domains[11]. Thus, the inter domain interactions are attributed to the high kinetic barriers in the early stages of unfolding[24, 25]. However, the origin of hysteresis is yet to be explored.

Experiments[10, 26, 27] further revealed a competitive protein aggregation pathway during the refolding of H*γ*D and H*γ*C Crys at [GdmCl] < 1 M due to population of partially folded intermediates[10]. Domain swapping[28, 29, 30, 31, 32, 33] plays an important role in the aggregation of globular proteins such as prion proteins[34], cystatin C[35, 36], c-Src SH3[37], *β*2-microglobulin[38] etc., where it is believed to drive the early stages of aggregation. In proteins such as RNase A, which can swap secondary structures from both Ntd and Ctd, double domain swapping is observed leading to a variety of differently assembled oligomeric states[39, 40, 41, 42]. To elucidate the structure of aggregation prone intermediates and underlying aggregation mechanism, several studies have investigated the dimer formation in H*γ*D Crys, which is the simplest oligomer[14, 12, 43]. A transient intermediate state with an extruded N terminal *β* hairpin loop is identified in single molecule atomic force microscopy (AFM) experiments[12] and computer simulations[43], which undergoes dimerization yielding different dimer structures in experiments and simulations. Although the molecular events leading to aggregation are not fully understood, domain swapping is considered to be the most probable aggregation mechanism[14, 12, 43] in H*γ*D Crys.

All-atom simulations predicted domain swapped structures where Ntd of one monomer interacts with the three *β* strands from *M*_4_ of Ctd from another monomer[14], and *β*_1_ from extruded *β* hairpin loop in Ntd forms antiparallel hydrogen bonds with the second *β* strand from *M*_4_ of Ctd [43]. These structures remain to be validated experimentally. AFM experiments on H*γ*D Crys indicated a different domain swapped structure with Ntd *β* hairpin loops swapped between adjacent monomers[12]. Experiments also suggest that the aggregation prone intermediate is not structurally similar to the unfolding intermediate and consists of a partially folded Ctd[44].

To probe the reasons behind the observed hysteresis during the folding of *γ* Crys and to gain insight into the aggregation prone misfolded intermediates, we studied the folding thermodynamics of H*γ*C (protein data bank (PDB) ID: 2NBR)[45] and H*γ*D (PDB ID: 1HK0)[46] Crys using molecular dynamics simulations and coarse grained self-organized polymer side-chain (SOP-SC) protein model[47, 48]. The folding free energy surface (FES) constructed from low friction Langevin dynamics simulations show that H*γ*C and H*γ*D Crys exhibit kinetic partitioning mechanism (KPM) and folds through at least three distinct pathways populating different intermediate states. One of the intermediates is an early unfolding intermediate with both domains folded and an exposed interface. Through the population of this intermediate, we show that the native proteins form a domain swapped dimer, which is a more stable entity with four folded domains and two interfaces. The population of this dimer structure during the early stages of unfolding can lead to hysteresis by resisting unfolding and requiring relatively higher denaturant concentrations to unfold. Experiments report similar domain swapped dimer structures for human *β*B2 Crys[49, 50, 51] (PDB ID: 2BB2) and mouse *γ*S Crys (PDB ID: 6MYG, 6MYH).

Experiments[10] further show that *γ* Crys exhibits significant aggregation while refolding at lower denaturant concentrations. To gain insight into this *in vivo* aggregation, we carried out dimerization simulations of the unfolded proteins. During dimerization, secondary structural elements from partially folded Ntd and Ctd of the proteins took part in domain swapping leading to an ensemble of double domain swapped structures. The primary swapping units in H*γ*C and H*γ*D Crys were found to be very similar, which can be attributed to their high sequence similarity (≈ 71 %). Both in H*γ*D and H*γ*C Crys, Ntds undergo aggregation by swapping the *β*_1_*β*_2_ hairpin loop, while Ctds swap the first hairpin loop in motif *M*_3_ (*β*_6_*β*_7_ in H*γ*D Crys, *β*_8_*β*_9_ in H*γ*C Crys). Other less common swapping units identified from the dimer structures include strands *β*_1_, *β*_3_*β*_4_, motif *M*_1_ in the Ntd of H*γ*D Crys (strands *β*_1_, *β*_7_ and motif *M*_1_ in the Ntd of H*γ*C Crys) and strands *β*_6_, *β*_12_ in the Ctd of H*γ*D Crys (*β*_8_, *β*_14_ in the Ctd of H*γ*C Crys). AFM experiments[12] on H*γ*D Crys also predicted a domain swapped dimer structure where N-terminal *β* hairpin loops get swapped between the monomers. Interestingly, a large number of cataract associated mutations are found in *β*_1_*β*_2_ loop region in H*γ*D Crys[24, 52, 53, 54, 55]. Such double domain swapping mechanism is feasible in an end to end fashion[56] among specific partially structured intermediates, giving rise to long chains of interconnected oligomers, which might act as aggregation precursors in cataractous lenses.

## Results

### Folding Thermodynamics of H*γ*C and H*γ*D Crys

Folding of *γ* Crys proteins have been studied using different experimental methods including fluorescence spectroscopy, circular dichroism, AFM etc[12, 44, 10, 22, 11]. However, detailed characterization of the transient intermediates populated in the folding pathways still remains a challenge. We performed low friction Langevin dynamics simulations to study the folding of H*γ*C and H*γ*D Crys using coarse grained SOP-SC protein model[47, 48]. A detailed description of the SOP-SC protein model and simulation parameters are in the supporting information (SI). The simulations are performed at temperatures ranging from *T* = 300 K to 500 K and we computed various thermodynamic properties associated with the folding.

Temperature dependent variation in specific heat capacity (*C*_*v*_) and average potential energy (⟨*E*⟩) indicate a cooperative two-state folding for H*γ*D Crys with the melting temperature, *T*_*M*_ ≈ 426 K (Figure 1C). Whereas a three-state folding with the population of an intermediate is observed for H*γ*C Crys with two peaks in the *C*_*v*_ plot at 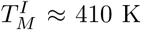 and 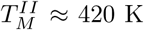 (Figure 2B). Experimentally[57, 26] measured *T*_*M*_ for H*γ*C and H*γ*D Crys are 352 K and 357 K, respectively. The deviation in the *T*_*M*_ obtained from experiments and simulations is due to the coarse-grained level description used to model the proteins, which cannot accurately mimic the experimental conditions. Experimental studies on H*γ*D and H*γ*C Crys further indicated a three state equilibrium unfolding with the population of an intermediate state[10]. Although the *C*_*v*_ plot associated with H*γ*C Crys indicates a three state folding/unfolding transition with an intermediate, only two state folding/unfolding transition is visible from the *C*_*v*_ plot of H*γ*D Crys.

**Figure 2:**
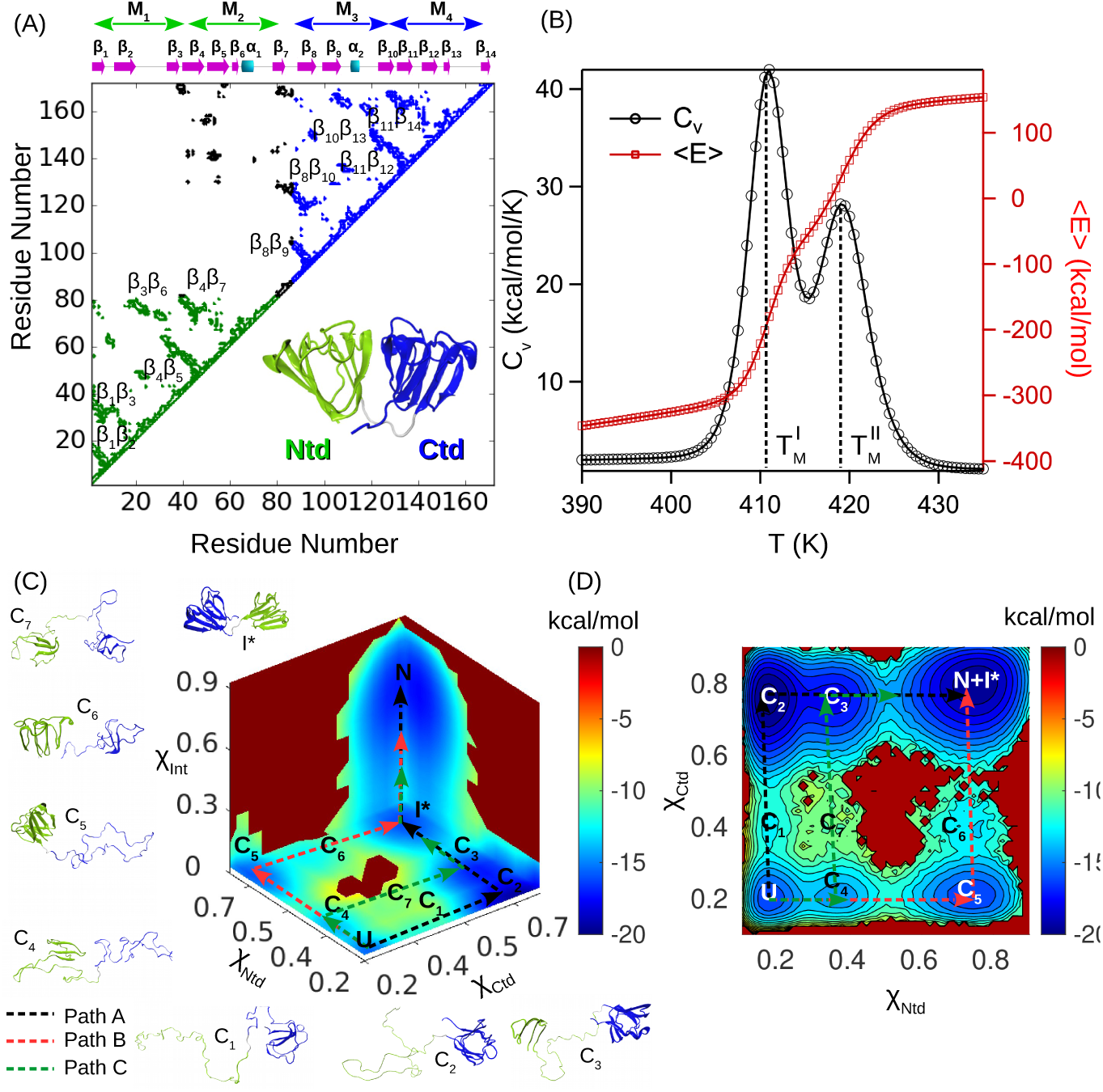
(A) Contact map of H*γ*C Crys[46] (PDB ID: 2NBR)[45]. H*γ*C Crys consists of two domains called N-terminal domain (Ntd, shown in green) and C-terminal domain (Ctd, shown in blue) connected by a short peptide linker. Each domain is further composed of two intercalated greek key motifs. Greek key motifs constituting the Ntd are termed as *M*_1_ (residues Gly1 to Ser39) and *M*_2_ (residues Gly40 to Pro82), while motifs constituting the Ctd are termed as *M*_3_ (residues His87 to Glu127) and *M*_4_ (residues Gly128 to Val170). The intra domain contacts are colored green (Ntd) and blue (Ctd), while inter domain contacts are colored black. (B) Temperature dependent variation in *Cv* and *(E)* indicate a three state folding transition in H*γ*C Crys. (C) *F* (*χ*_*Ntd*_, *χ*_*Ctd*_, *χ*_*Int*_) projected onto *χ*_*Ntd*_, *χ*_*Ctd*_ and *χ*_*Int*_ at 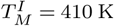. Three pathways connect the *N* state and *U* state through distinct intermediate states. Representative structures of the intermediates are shown. (D) *F* (*χ*_*Ntd*_, *χ*_*Ctd*_) projected onto *χ*_*Ntd*_ and *χ*_*Ctd*_ at 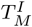. All the intermediate states are visible except for *I*^∗^ (Ctd and Ntd domains are folded with disrupted inter domain contacts), which is indistinguishable from the *N* state.

We also estimated the *T*_*M*_ of Ntd and Ctd domains of both the proteins from the full-length protein simulations by computing the variance in the fraction of native contacts in the Ntd (*χ*_*Ntd*_) and Ctd domains (*χ*_*Ctd*_) as a function of *T* (see SI for details). The *T*_*M*_ of Ntd and Ctd domains approximately corresponds to the peak position in the plots (Figure S1, S2). The Ntd and Ctd of H*γ*C Crys exhibit differential domain stability with *T*_*M*_ of Ntd and Ctd ≈ 410 K and ≈ 420 K, respectively. The *T*_*M*_ of Ntd and Ctd domains corresponds to 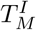 and 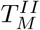 of H*γ*C Crys in the *C*_*v*_ plot (Figure 2B), indicating sequential unfolding of Ntd followed by Ctd. Thus the intermediate populated in the *C*_*v*_ plot (Figure 2B) of full-length H*γ*C Crys consists of a folded Ctd and an unfolded Ntd, evident from the *T*_*M*_ values of the individual domains. Unlike H*γ*C Crys, truncated Ntd and Ctd of H*γ*D Crys indicated comparable thermal stabilities (with *T*_*M*_ ≈ 426 K) in agreement with the *C*_*v*_ plot of full-length H*γ*D Crys (Figure 1C). However, experiments[27] report a higher thermodynamic stability of the Ctd (*T*_*M*_ = 349.2 K) as compared to the Ntd (*T*_*M*_ = 337.5 K) of H*γ*D Crys. This anomaly arises due to the reduced-level description of the model systems, as discussed earlier.

### FES of H*γ*D and H*γ*C Crys Folding Show Population of Multiple Intermediate States

Although thermodynamic properties such as ⟨*E*⟩ and *C*_*v*_ capture major transitions associated with protein folding/unfolding, they often fail to detect transient intermediates, which occur due to minor structural perturbations that play an important role in protein aggregation. To identify the transient intermediates we constructed the FES associated with the folding of H*γ*D and H*γ*C Crys. The FES is projected onto 3 collective variables (CVs): (1) fraction of native contacts in Ntd (*χ*_*Ntd*_), (2) fraction of native contacts in Ctd (*χ*_*Ctd*_) and (3) fraction of native contacts at the interface of Ntd and Ctd (*χ*_*Int*_) (see Methods for details). We employed weighted histogram analysis method (WHAM)[58] to construct the three-dimensional FES, *F* (*χ*_*Ntd*_, *χ*_*Ctd*_, *χ*_*Int*_).

The folding FES of H*γ*D and H*γ*C Crys show that for both the proteins, at least 3 distinct minimum energy pathways connect the native (*N*) and unfolded (*U*) basins populating different intermediates (Figure 1D,E,2C,D,S3,S4). In H*γ*D Crys, the most preferred pathway (Path A, shown in black) corresponds to the folding of the Ctd followed by the folding of Ntd (Figure 1D). In this pathway, 4 intermediates are populated: (1) a partially folded Ctd with motif *M*_4_ folded and Ntd unfolded (basin *D*_1_), (2) Ctd completely folded and Ntd unfolded (basin *D*_2_), (3) Ctd completely folded and motif *M*_1_ of Ntd folded (basin *D*_3_), and (4) both Ctd and Ntd folded with disrupted inter-domain contacts (basin *I*^∗^) (Figure S3A,B,C,H,S5A). Intermediate *D*_2_ with a folded Ctd and unfolded Ntd was inferred from experiments[22, 9] and was also observed in the H*γ*D unfolding simulations[13, 15].

In the second preferred pathway (Path B, shown in red), Ntd folds prior to the folding of Ctd (Figure 1D). In this pathway, 4 intermediates are populated: (1) a partially folded Ntd with motif *M*_1_ folded and Ctd unfolded (basin *D*_4_), (2) completely folded Ntd and unfolded Ctd (basin *D*_5_), (3) completely folded Ntd and partially folded Ctd with motif *M*_4_ folded (basin *D*_6_) and (4) intermediate *I*^∗^, which is also common to Path A (Figure S3D,E,F). AFM experiments on H*γ*D Crys provide evidence for the population of an intermediate with a folded Ntd and unfolded Ctd, which is similar to the intermediate, *D*_5_. Interestingly, kinetic unfolding experiments on H*γ*D Crys in the absence of denaturants indicated higher kinetic stability of Ntd compared to Ctd[11].

In Paths A and B, sequential folding of the domains is observed. Whereas in the third folding pathway (Path C, shown in green), which is the least favored, intermediate with both the domains partially folded are observed. An intermediate with partially folded Ntd with motif *M*_1_ folded and partially folded Ctd with motif *M*_4_ folded (basin *D*_7_) is populated in addition to *D*_4_, *D*_3_ and *I*^∗^ (Figure S3G,S5A).

Similar to H*γ*D Crys, the preferred path connecting the *N* and *U* basins in H*γ*C Crys (Path A shown in black) involves stepwise folding of the Ctd prior to the folding of Ntd. The 4 intermediates populated in this path are: (1) partially folded Ctd with motif *M*_4_ folded and unfolded Ntd (basin *C*_1_), (2) completely folded Ctd and unfolded Ntd (basin *C*_2_), (3) completely folded Ctd and partially folded Ntd with *M*_1_ folded (basin *C*_3_), and (4) both Ctd and Ntd folded with disrupted inter-domain contacts (*I*^∗^) (Figure 2C,S4A,B,C,H,S5B). In the second preferred path (Path B shown in red), Ntd folds prior to Ctd. The 4 intermediates populated in this path are (1) partially folded Ntd with folded motif *M*_1_ and unfolded Ctd (basin *C*_4_), (2) completely folded Ntd and unfolded Ctd (basin *C*_5_), (3) completely folded Ntd and partially folded Ctd with motif *M*_4_ folded (basin *C*_6_), and (4) intermediate *I*^∗^ (Figure S4D,E,F,S5B). The least favored path (Path C shown in green) involves an intermediate with partially structured Ntd with motif *M*_1_ folded and partially structured Ctd with motif *M*_4_ folded (basin *C*_7_) in addition to *C*_3_, *C*_4_ and *I*^∗^ (Figure S4G,S5B).

The intermediate, *I*^∗^, which is common to all the folding pathways of H*γ*D and H*γ*C Crys is populated due to structural fluctuations in the native state ensemble. A hydrophobic patch at the domain interface of Ntd and Ctd gets exposed when *I*^∗^ is populated and this makes it an ideal candidate for domain swapping. Human *β*B_2_ Crys, which belongs to the *βγ* Crys superfamily, forms such domain swapped dimers by swapping the Ntd and Ctd while the linker region acts as the hinge loop[51]. Similar domain swapped oligomers are also found in mouse *γ*S Crys, which is a member of *γ* Crys family. Although no such domain swapped structures are reported for H*γ*D and H*γ*C Crys, intermediate *I*^∗^ can potentially lead to such domain swapped oligomers. A recent computational study on W42R mutant of H*γ*D Crys showed the presence of *I*^∗^ intermediates with exposed interdomain hydrophobic interface, which led to large scale aggregation involving Ntd-Ntd interactions[59].

Rest of the partially structured intermediate states populated in the folding pathways of H*γ*D and H*γ*C Crys can act as potential precursors to aggregation as well. Formation of sticky hydrophobic patches on the protein surface has been related to aggregation in cataract associated mutants of H*γ*D Crys[53, 60, 61]. Experimental studies on H*γ*D Crys indicated the presence of intermediates with either Ctd folded (basin *D*_2_)[22, 9] or Ntd folded (basin *D*_5_)[12, 11]. However, the role of these intermediates in aggregation remains unclear. Domain swapped dimer structure of H*γ*D Crys was predicted in AFM experiments, where *β*_1_*β*_2_ strands from Ntd get swapped between neighboring monomers[12]. Further, fluorescence spectra of chaperone (*α*B Crys) bound complex of H*γ*D Crys showed that both the domains were partially unstructured in the substrate[44], unlike the experimentally observed folding intermediates, which consist of a folded domain. Intermediates such as *D*_7_, *C*_7_ populated in the FES of H*γ*D and H*γ*C Crys, consist of partially unfolded Ntd and Ctd, and meet the requirements for possible aggregtaion prone precursors. These intermediates or structurally related intermediates with exposed hydrophobic interface might play an important role in the aggregation of *γ* Crys proteins.

### Dimerization in H*γ*D and H*γ*C Crys

Surgically removed cataractous lenses contain high molecular weight aggregates of partially folded *γ* Crys[62, 26, 44]. However, the specific conformations of the proteins in the aggregates remain elusive due to the insoluble nature of the aggregates. A detailed understanding of the misfolded conformations is crucial to elucidate the underlying molecular mechanism of cataract formation and design essential therapeutics. To understand the initial molecular interactions that advances to cataract formation in later stages, we studied the formation of dimers, which are the simplest possible oligomers, using symmetric Go potential to model inter-monomer interactions[32, 63] (Figure S9).

### Dimerization from Native State

*γ* Crys are generally monomeric in solution. However, octameric structures of mouse *γ*S Crys are published in the PDB, where Ntds and Ctds are swapped between the neighboring monomers with linker region acting as hinge loop (PDB ID: 6MYG, 6MYH), indicating oligomerization in the native state of the protein. Similar domain swapping is also observed in human *β*B_2_ Crys[51], which belongs to the *βγ* Crys superfamily. To elucidate whether such oligomerization is also feasible in the native states of H*γ*D and H*γ*C Crys, we performed 50 independent Brownian dynamics simulations starting from two folded monomers (say monomers A and B) separated by 20 Å (≈ *R*_*g*_ of a monomer) using a harmonic restraint with a spring constant, *k* = 2.0 kcal/mol/Å^2^ at *T* = 300 K. To track the progress of dimerization, we computed the number of contacts *Q*_*AB*_ formed between the monomers A and B as a function of simulation time (Figure 3A,B). A two step dimerization process was observed in both the proteins yielding unique dimer structures (*Dim*_*γD*_ and *Dim*_*γC*_).

**Figure 3:**
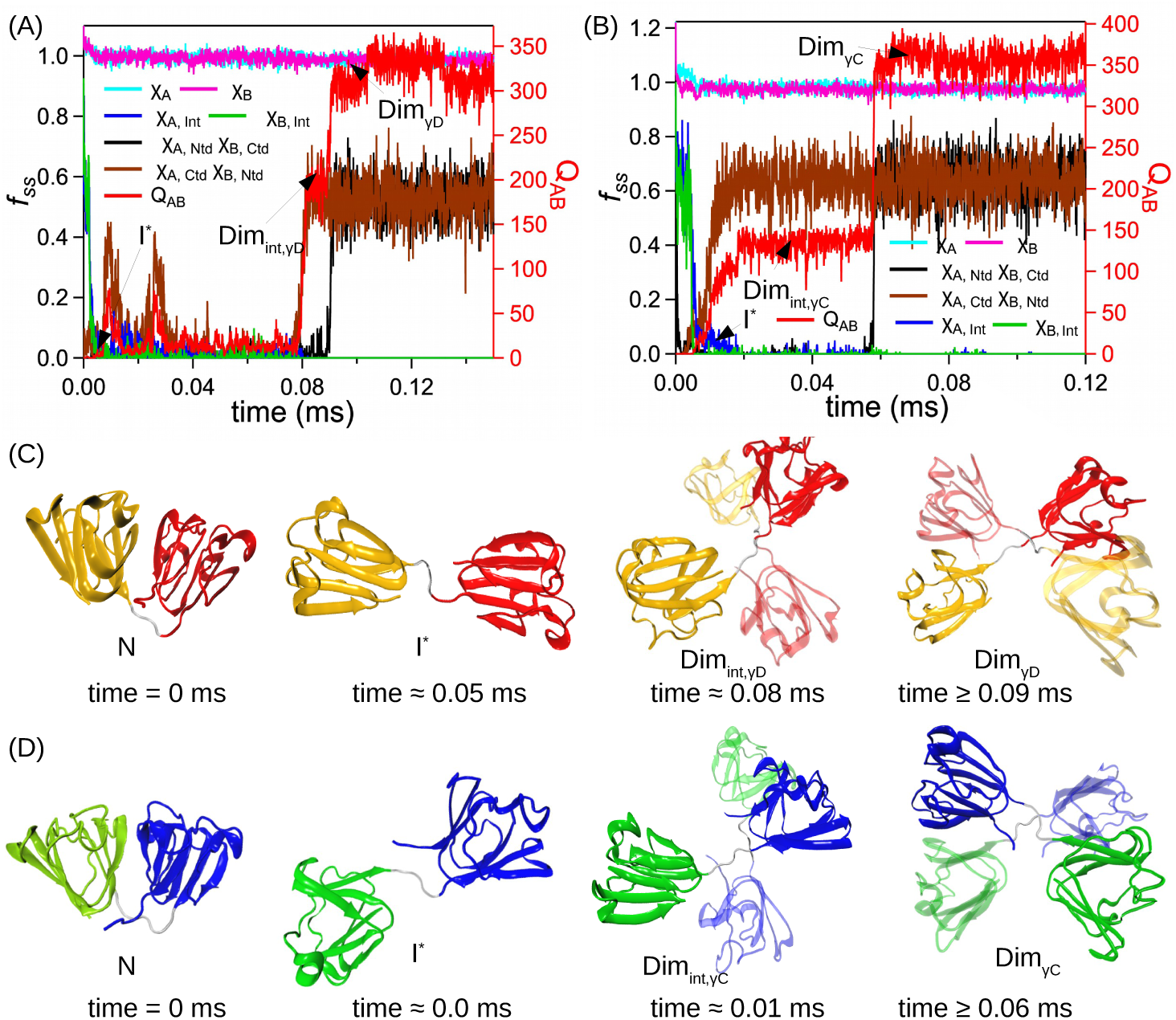
Dimerization from the native states of H*γ*D and H*γ*C Crys. (A, B) Time evolution of the fraction of secondary structural contacts (*fss*) present in the two monomers A and B, and the total number of native contacts *Q*_*AB*_ formed between A and B. Dimerization takes place in two steps. In the first step, only one domain gets swapped (*Dim*_*int,γD*_, *Dim*_*int,γC*_) while in the second step both the domains are swapped (*Dim*_*γD*_, *Dim*_*γC*_). Mechanism of dimerization in H*γ*D (C) and H*γ*C Crys (D). An intermediate state *I*^∗^ with disrupted inter domain contacts get populated in the initial stages of dimerization. *I*^∗^ further swaps the Ntd and Ctd with neighboring monomers in two stages leading to domain swapped dimers.

In the final dimer structure, both Ntd and Ctd of the two monomers remain folded (fraction of native contacts in monomer A, *χ*_*A*_ ≈ 1 and in monomer B, *χ*_*B*_ ≈ 1), however the interdomain contacts present between Ntd and Ctd of each monomer get disrupted (*χ*_*A,Int*_ ≈ 0, *χ*_*B,Int*_ ≈ 0) (Figure 3A,B). Instead, two new interfaces form between Ntd of monomer A and Ctd of monomer B (*χ*_*A,Ntd*;*B,Ctd*_ ≈ 1), and Ctd of monomer A and Ntd of monomer B (*χ*_*A,Ctd*;*B,Ntd*_ ≈ 1). This reveals a domain swapping mechanism, where neighboring monomers exchange Ntd and Ctd with each other forming two new interfaces, while the linker region acts as hinge loop. Dimers *Dim*_*γD*_ and *Dim*_*γC*_ obtained for H*γ*D and H*γ*C Crys, respectively, are very similar to the domain swapped structure obtained for mouse *γ*S Crys (PDB ID: 6MYG, 6MYH). NMR studies[64] further show that amorphous aggregates of cataract associated P23T mutant of H*γ*D Crys retains the native fold, indicating native-like conformations taking part in aggregation.

In the first step of dimerization, proteins break their inter domain contacts exposing the hydrophobic pocket present at the interface, populating the intermediate *I*^∗^. In *I*^∗^, the linker region adopts an extended conformation, which facilitates domain swapping (Figure 3C,D). In the next step, monomers swap either Ntd or Ctd with each other leading to half swapped dimers *Dim*_*int,γD*_ and *Dim*_*int,γC*_. Finally the remaining domains are swapped between the monomers resulting in fully swapped dimers *Dim*_*γD*_ and *Dim*_*γC*_. Intermediate *I*^∗^, which is key to dimerization in native state gets populated during the initial stages of unfolding of both the proteins (Figure 1D,2C). Low denaturant concentrations, which introduce small scale perturbations in the native structure, can enhance the population of *I*^∗^. *I*^∗^ states can further take part in domain swapping with neighboring monomers leading to long chain of oligomeric structures. Population of such dimeric or oligomeric moities with multiple inter protein interfaces can resist unfolding to a relatively higher denaturant concentrations and lead to hysteresis as observed in experiments[10, 23, 24].

### Dimerization During Refolding

Aggregation is also observed experimentally for H*γ*D and H*γ*C Crys when [GdHCl] is lowered from high values to less than 1 M[10]. Aggregation suppression experiments on *γ* Crys using chaperone *α*B Crys further identified partially unfolded substrates of H*γ*C, H*γ*D and H*γ*S Crys in the aggregates[44]. It was further concluded that both Ntd and

Ctd of H*γ*D Crys must be partially unfolded for *in-vitro* aggregation[44]. To identify conformations of H*γ*D and H*γ*C Crys populated during refolding that result in aggregation, we studied dimerization of the proteins during refolding from unfolded conformations at *T* = 300 K. We performed 100 independent Brownian dynamics simulations starting from different unfolded conformations of the monomers separated by ≈ 20 Å. Dimerization was observed in ≈ 60% of the trajectories in H*γ*D and H*γ*C Crys.

An ensemble of domain swapped dimer structures were identified, where at least in 45% of the structures of H*γ*D Crys (≈ 40% of H*γ*C Crys), secondary structural units were swapped from both Ntd and Ctd resulting in double domain swapped dimers. In rest of the structures, either Ntd (≈ 40% in H*γ*D Crys, ≈ 37% in H*γ*C Crys) or Ctd (≈ 15% in H*γ*D Crys, ≈ 23% in H*γ*C Crys) took part in swapping, while the other domain folded to its native structure. In both the proteins, most populated double domain swapped dimers are formed by swapping the secondary structural units from both Ntd and Ctd. The primary swapping element in the Ntds is the first hairpin loop *β*_1_*β*_2_ from motif *M*_1_ (Figure 4A,C, S6A,C). Swapping of *β*_1_*β*_2_ strands was predicted in an AFM study of H*γ*D Crys[12]. In Ctds (Figure 4B,C, S6B,C), the primary swapping units are the first hairpin loop of motif *M*_3_ (*β*_6_*β*_7_ in H*γ*D Crys and *β*_8_*β*_9_ in H*γ*C Crys).

**Figure 4:**
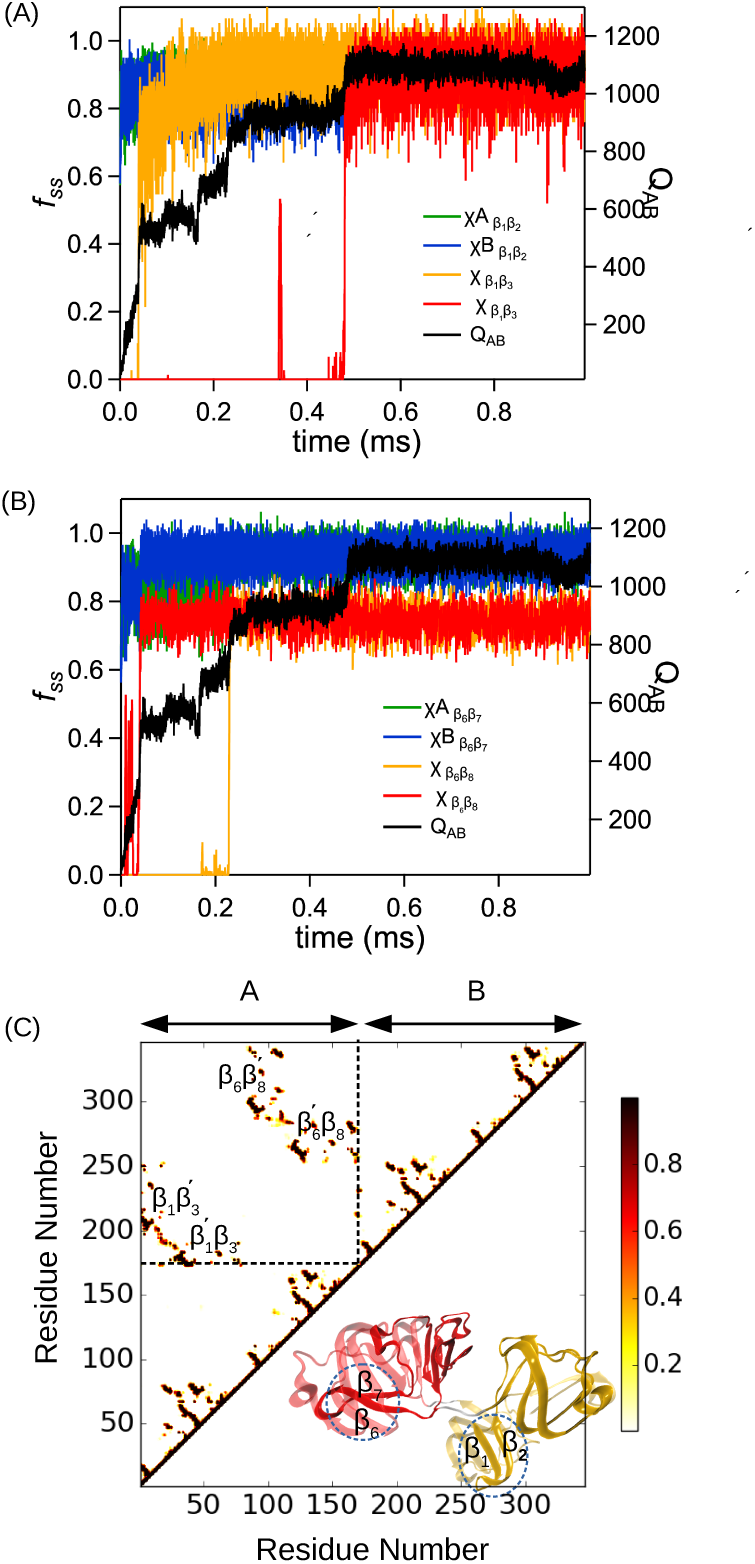
Multistep dimerization in H*γ*D Crys while refolding at *T* = 300 K. Secondary structural elements from Ntd (*β*_1_*β*_2_) and Ctd (*β*_6_*β*_7_) get swapped leading to a double domain swapped dimer. (A) Time evolution of fraction of secondary structural contacts between *β*_1_*β*_2_ strands in monomer 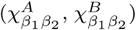 and between *β*_1_ strand of monomer A and 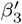 strand of monomer 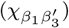 and vice versa 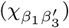 along with the number of contacts present between monomer A and B (*Q*_*AB*_). (B) Time evolution of fraction of secondary structural contacts between *β*_6_*β*_7_ strands in monomer A and 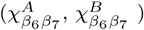 and between *β*_6_ strand of monomer A and 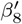 strand of monomer 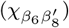 and vice versa 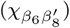. (C) Contact map of the final domain swapped dimer structure. *β*_1_ strand makes contacts with 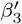 strand of the other monomer and vice versa after swapping of *β*_1_*β*_2_ strands. The first hairpin loop of *M*_3_ (*β*_6_*β*_7_) get swapped between the Ctds where *β*_6_ strand forms contact with 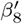 strand from the other monomer and vice versa.

In ≈ 10% of the cases, we observed domain swapped structures similar to *Dim*_*γD*_, *Dim*_*γC*_ where both the domains are completely swapped between the monomers. Other less common swapping units (with population < 5 %) were identified in H*γ*D and H*γ*C Crys, which led to both single and double domain swapped dimer structures (Figure S7,S8). Ntd swapping units include *β*_1_, *β*_3_*β*_4_, motif *M*_1_ in H*γ*D Crys, and *β*_1_, *β*_7_, motif *M*_1_ in H*γ*C Crys. Ctd swapping units include *β*_6_, *β*_12_ in H*γ*D Crys, and *β*_8_, *β*_14_ in H*γ*C Crys. Some of the intermediates populated in the folding FES of H*γ*D (*D*_3_, *D*_4_, *D*_7_) and H*γ*C Crys (*C*_3_, *C*_4_, *C*_7_) with a folded motf *M*_1_ should facilitate swapping of *M*_1_ between the Ntds as observed in the ensemble of swapped dimer structures.

In the most populated dimers, domain swapping took place after formation of the *β* hairpin loops *β*_1_*β*_2_, *β*_6_*β*_7_ (in H*γ*D Crys) and *β*_8_*β*_9_ (in H*γ*C Crys) (Figure 4, S6). In the final domain swapped structures, *β*_1_ strand makes contacts with 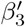 strands from the other monomer and vice versa. While in Ctds, *β*_6_ strand in H*γ*D Crys (*β*_8_ strand in H*γ*C Crys) makes contacts with 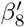 strand (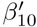 strand in H*γ*C Crys) from the other monomer and vice versa. Interestingly, in H*γ*D Crys, highest number of pathogenic mutations are located in *β*_1_*β*_2_ hairpin loop[24, 52, 53, 54, 55]. The double domain swapping mechanism reported in this study can be used to explain higher order oligomer formation in *γ* Crys where instead of reciprocal domain swapping, secondary structures can be exchanged with neighboring monomers in a chain-like manner.

## Conclusions and Outlook

Cataract associated misfolding and aggregation mechanism of eye lens Crys proteins remains largely unknown. Using coarse-grained models and molecular dynamics simulations, we explored the folding pathway and dimerization of H*γ*D and H*γ*C Crys. The multi-dimensional FES shows three competing folding pathways with distinct intermediates. In the two dominant pathways, the domains sequentially fold, populating intermediates with either Ntd or Ctd folded. The third, least favored pathway, populates intermediates with partially structured Ntd and Ctd, which can facilitate aggregation from both the domains. Interestingly, fluorescence spectra of chaperone (*α*B Crys) bound complex of H*γ*D Crys indicated that both the domains are partially structured in the aggregation prone conformations[44]. All three pathways populate a native-like intermediate state (*I*^∗^), with two folded domains and disrupted inter-domain contacts. Under native-like conditions, *I*^∗^ forms domain swapped dimers by exchanging folded domains with other monomers. Such oligomerization, if takes place during early stages of unfolding, may lead to hysteresis in the folding experiments. Similar domain swapped structures are reported for human *β*B2 Crys[49, 50, 51] and mouse *γ*S Crys.

We have also identified an ensemble of domain swapped dimers from the dimerization simulations performed using unfolded conformations of H*γ*D and H*γ*C Crys, which mimic aggregation during refolding. The Ntd swapping units (*β*_1_*β*_2_) for H*γ*D Crys were already hinted in previous studies[12, 43, 65] and contains the highest number of pathogenic mutations. We also identified swapping units in the Ctds (*β*_6_*β*_7_ in H*γ*D and *β*_8_*β*_9_ in H*γ*C Crys), which in conjunction with Ntd swapping units led to double domain swapped dimers. Other less populated dimers involved swapping of different secondary structures from the Ntds and the Ctds, leading to heterogeneity in the dimer structures. The double domain swapping mechanism presented in this study provides structural insights to the early oligomeric assembies in H*γ*D and H*γ*C Crys. Knowledge of the swapping units present in the two domains can be exploited to design preventive measures and essential therapeutics for cataract.

The coarse-grained model used in this study has been successfully applied to elucidate folding[48, 66, 67] and aggregation of other proteins[68, 69, 70]. However, the native-centric nature of this model limits the incorporation of non-native interactions that might play a crucial role in protein aggregation. Due to the low resolution of coarse-grained model, atom level description of protein-protein and solvent-protein interactions are also beyond the scope of this model. In the SOP-SC model used in this work, we have not incorporated features that allow the formation of non-native disulfide bonds. However, such disulfide bonds have been proven to be very important in the formation of crystallin aggregates.

Age related nuclear cataract often results from the oxidative stress built up in aged lens due to gradual depletion in the glutathione (GSH) levels[71, 72]. Due to oxidative stress, Cys residues form disulfide links, which are found commonly in mature cataractous lenses[73, 74, 75, 76]. Consequently, impact of disulfide bonds in aggregation of Crys proteins, especially in Cys rich *γ* Crys is the subject of intense investigation[77, 78, 79, 43, 65]. A recent experiment[65] provided evidence for dynamic transfer of disulfide bonds from oxidized WT H*γ*D Crys to a mutant W42Q promoting aggregation in the latter. Another experimental study[43] in conjunction with simulations identified an internal non-native disulfide bond (Cys32-Cys41), which traps an aggregation prone conformation, essential for oxidative aggregation of mutant variants (W42R, W42Q) of H*γ*D Crys. The trapped intermediate obtained from the mutant simulations, features a detached N-terminal *β*-hairpin. This intermediate is structurally similar to the intermediate predicted in AFM experiments[12] of WT H*γ*D Crys and also consistent with the domain swapped dimer structures predicted in this work for the WT protein. This shows that aggregation prone intermediates, which are predominantly populated by mutants, are transiently populated in WT proteins as well. In our future computational studies, we will probe the role of non-native disulfide bonds in the population of aggregation-prone structures by *γ* Crys proteins using coarse-grained protein models[80, 81], which allow the formation of such bonds.

## Materials and Methods

Molecular dynamics simulations are performed using coarse-grained self-organized polymer side-chain (SOP-SC)[47, 48] description of the human *γ* Crys proteins. Coarse-grained structures of the proteins are built from their PDB structures with PDB IDs 2NBR[45] (H*γ*C Crys) and 1HK0[46] (H*γ*D Crys). The Hamiltonian of a protein conformation in the SOP-SC model represented by a set of coordinates {**r**} is,

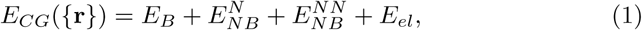

where *E*_*B*_ is the energy due to bonded interaction present between the covalently bonded beads, 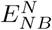 and 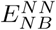 are the energy due to non-bonded native and non-native interactions, respectively, and *E*_*el*_ is the energy due to electrostatic interactions present between the charged residues in the protein. Native interactions exist between two beads, if they are separated by at least three bonds and are within a cut-off distance (*R*_*c*_ = 8 Å) in the SOP-SC model of the PDB structure while any other non-covalent interactions are considered as non-native. Symmetric Go potential[32, 63] (Figure S9) is used to model the interactions between two protein monomers which is discussed in details in the supporting information (SI). Langevin dynamics simulations are used to compute equilibrium thermodynamic properties. Weighted histogram analysis method is used to construct the three-dimensional free energy landscape *F*(*χ*_*Ntd*_, *χ*_*Ctd*_, *χ*_*Int*_) from the Langevin dynamics simulations performed at various temperatures. Brownian dynamics[82] simulations are performed to study dimerization kinetics of the proteins. A detailed description of the protein models, simulation methodolgy and parameters are provided in the SI.

## Acknowledgments

G.R. acknowledges funding from the National Supercomputing Mission (NSM) through the grant MeitY/R&D/HPC/2(1)/2014. J.N. thanks DST-SERB for funding under Ramanujan Faculty Fellowship. B.M. acknowledges research fellowship from Indian Institute of Science-Bangalore. The computations are performed using the TUE and Cray XC40 cluster at IISc.

