## Supporting Information for "Double Domain Swapping in Human *γ*C and *γ*D Crystallin Drives Early Stages of Aggregation"

### Supporting Information for "Double Domain Swapping in Human $\gamma$C and $\gamma$D Crystallin Drives Early Stages of Aggregation"

Balaka Mondal

*Solid State and Structural Chemistry Unit, Indian Institute of Science, Bangalore, Karnataka 560012, India*

Jayashree Nagesh

*Solid State and Structural Chemistry Unit, Indian Institute of Science, Bangalore, Karnataka 560012, India*

Govardhan Reddy*

*Solid State and Structural Chemistry Unit, Indian Institute of Science, Bangalore, Karnataka 560012, India*

*Corresponding author

*Preprint submitted to Journal of Molecular Biology* *July 19, 2020*

#### 1. Methods

**Self Organized Polymer-Side Chain (SOP-SC) Model for Proteins:** We have used native-centric SOP-SC model[1, 2] to study the folding thermodynamics of H$\gamma$C Crys and H$\gamma$D Crys. In SOP-SC model, each amino acid is represented by two beads. The backbone atoms of an amino acid are represented by a bead positioned at the center of $C_\alpha$ atom, and the side chain atoms are represented by another bead positioned at the center of mass of the side-chain. The SOP-SC model for H$\gamma$C Crys and H$\gamma$D Crys are constructed using the structures in the protein data bank (PDB) with PDB ID: 2NBR[3] and 1HK0[4], respectively. Missing hydrogen atoms are added to the structures using the program visual molecular dynamics (VMD) [5] before calculating the centre of mass of the side-chain. The Hamiltonian corresponding to the SOP-SC model is described in terms of bonded ($E_B$), non-bonded ($E_{NB}$) and electrostatics interactions ($E_{el}$). Covalently connected beads interact via a bonded potential ($E_B$). Non-bonded interactions ($E_{NB}$) consist of native (N) and non-native (NN) interactions. Interactions between two beads are considered native, if they are separated by at least three bonds and are within a cut-off distance ($R_c$) in the SOP-SC model of the PDB structure. Any other non-covalent interactions are considered as non-native interactions ($E_{NN}$). Native interactions between neighboring side-chain beads are ignored because of their close proximity in both folded and unfolded states. Electrostatic interactions are present between charged residues and are ignored if charges are present on adjacent side-chain beads. The force-field associated with the SOP-SC model for a protein conformation described by the set of coordinates $\{\mathbf{r}\}$ is given by,

$$E_{CG}(\{\mathbf{r}\}) = E_B + E_{NB}^{N} + E_{NB}^{NN} + E_{el} \tag{S1}$$

The bonds between the beads in the SOP-SC model are modeled using the finite extensible nonlinear elastic (FENE) potential and is given by,

$$E_B = -\sum_{i=1}^{N_B} \frac{k}{2} R_0^2 \log\Big(1 - \frac{(r_i - r_{cry,i})^2}{R_0^2}\Big), \tag{S2}$$

$N_B$ being the total number of bonds present between the covalently linked beads, $r_i$ is the distance between $i^{th}$ pair of beads and $r_{cry,i}$ is the distance between the same $i^{th}$ pair of beads in the SOP-SC PDB structure. The native interactions $E^N_{NB}$, are modeled using a Lennard-Jones type of potential given as,

$$\begin{aligned} E^N_{NB} = &\sum_{i=1}^{N_N^{bb}} \epsilon_h^{bb} \Big[ \big(\frac{r_{cry,i}}{r_i}\big)^{12} - 2\big(\frac{r_{cry,i}}{r_i}\big)^6 \Big] + \sum_{i=1}^{N_N^{bs}} \epsilon_h^{bs} \Big[ \big(\frac{r_{cry,i}}{r_i}\big)^{12} - 2\big(\frac{r_{cry,i}}{r_i}\big)^6 \Big] \\ &+ \sum_{i=1}^{N_N^{ss}} 0.5 \times 300 k_B \times (0.7 - \epsilon_i^{ss}) \Big[ \big(\frac{r_{cry,i}}{r_i}\big)^{12} - 2\big(\frac{r_{cry,i}}{r_i}\big)^6 \Big], \end{aligned} \tag{S3}$$

where $N_N^{bb}$, $N_N^{bs}$ and $N_N^{ss}$ represent the total number of native contact pairs present between backbone-backbone, backbone-side chain, and side chain-side chain beads, respectively. $k_B$ is the Boltzmann constant, $r_i$ is the distance between $i^{th}$ pair of beads and $r_{cry,i}$ is the distance between the same $i^{th}$ pair of beads in the SOP-SC PDB structure. $\epsilon_h^{bb}$ and $\epsilon_h^{bs}$ denote the strength of interaction between backbone beads and backbone-side chain beads respectively. Strength of interactions between side-chain beads $\epsilon_i^{ss}$ are obtained from the Betancourt-Thirumalai statistical potential[6].

Purely repulsive non-native interactions ($E^{NN}_{NB}$) are modeled as

$$E^{NN}_{NB} = \sum_{i=1}^{N_{NN}} \epsilon_l \big(\frac{\sigma_i}{r_i}\big)^6 + \sum_{i=1}^{N_{ang}^{bb}} \epsilon_l \big(\frac{\sigma^{bb}}{r_i}\big)^6 + \sum_{i=1}^{N_{ang}^{bs}} \epsilon_l \big(\frac{\sigma_i^{bs}}{r_i}\big)^6, \tag{S4}$$

where $N_{NN}$ is the total number of non-native interaction pairs present in the SOP-SC model, $\sigma_i$ is sum of the radii of the beads in $i^{th}$ pair of non-native interactions, $\sigma^{bb}$ is the diameter of backbone bead, and $\sigma^{bs} = \frac{\sigma^{bb}+\sigma_i^{ss}}{2}$ where $\sigma_i^{ss}$ is the diameter of the side chain bead in $i^{th}$ angular interaction between a backbone and side chain beads. The second and third terms in eq. S4 model the bond angle potential between beads separated by two bonds. $N_{ang}^{bb}$ and $N_{ang}^{bs}$ represent the total number of bond angles between the backbone beads and between the backbone-side chain beads. Electrostatic interactions are implemented using a screened Coulomb potential given as

$$E_{el} = \sum_{i=1}^{N_c-1} \sum_{j=i+1}^{N_c} \frac{q_i q_j exp(-\kappa r_{ij})}{\epsilon r_{ij}}, \tag{S5}$$

where $N_c$ is the number of charged residues in the protein, $r_{ij}$ is the distance between the charged side chains $i$ and $j$, $q_i$ and $q_j$ are the point charges measured in units of electron charge placed on the centers of the charged side chain beads $i$ and $j$, respectively. At neutral pH, $q_i$ is considered +1 for positively charged residues, and -1 for negatively charged residues. The inverse Debye length, $\kappa$, accounts for the presence of a monovalent salt of 10mM concentration. For implicit solvent description, usual range of dielectric constants are from 2 to 20[7]. In the simulations, we considered the dielectric constant of the medium as $\epsilon = 10\ \epsilon_0$, where $\epsilon_0$ is the vacuum permittivity. Values of the parameters used to describe the SOP-SC energy functions and side-chain radii are listed in Table S1, Table S2 and Table S3, respectively.

**Symmetric Go potential to mimic inter-protein interactions:** To mimic the interactions between two protein chains, we employed symmetrized Go-type hamiltonians[8, 9], where native contacts present in a monomer chain is used to define the additional inter-protein interactions present in the dimeric system. For each native interaction present between residues $i$ and $j$ of a monomeric chain A, there are three additional native interactions present between identical residue $i'$ of another monomeric chain B and $j$ of chain A, $i$ of chain A and $j'$ of chain B and finally between $i'$ and $j'$ of chain B. The operative force field in the dimeric systems (monomer A and B) can be described as,

$$\begin{aligned} E_{A,B}(\{\mathbf{r}\}) = {} & E_B(A) + E_B(B) + E^{N}_{NB}(A) + E^{N}_{NB}(B) \\ & + E^{NN}_{NB}(A) + E^{NN}_{NB}(B) + E^{N}_{NB}(A,B) + E^{NN}_{NB}(A,B), \end{aligned} \tag{S6}$$

where $E_B(A)$ and $E_B(B)$ represent bonded interactions in the monomers A and B, $E^{N}_{NB}(A)$ and $E^{N}_{NB}(B)$ represent non-bonded native interactions in A and B, and $E^{NN}_{NB}(A)$ and $E^{NN}_{NB}(B)$ represent non-bonded non-native interactions in A and B. The terms $E^{N}_{NB}(A,B)$ and $E^{NN}_{NB}(A,B)$ in eq. S6 represent the inter native contacts and non-native contacts present between A and B, respectively.

The inter native interactions between the monomers A and B is given by

$$E^{N}_{NB}(A,B) = \sum_{i=1}^{N^{bb}_{N}(A,B)} \epsilon^{bb}_{h}\big[\big(\frac{r_{cry,i}}{r_i}\big)^{12} - 2\big(\frac{r_{cry,i}}{r_i}\big)^{6}\big] + \sum_{i=1}^{N^{bs}_{N}(A,B)} \epsilon^{bs}_{h}\big[\big(\frac{r_{cry,i}}{r_i}\big)^{12} - 2\big(\frac{r_{cry,i}}{r_i}\big)^{6}\big] + \sum_{i=1}^{N^{ss}_{N}(A,B)} 0.5\times 300k_B \times (0.7-\epsilon^{ss}_{i})\big[\big(\frac{r_{cry,i}}{r_i}\big)^{12} - 2\big(\frac{r_{cry,i}}{r_i}\big)^{6}\big], \tag{S7}$$

where $N^{bb}_{N}(A,B)$, $N^{bs}_{N}(A,B)$ and $N^{ss}_{N}(A,B)$ represent inter native backbone-backbone, backbone-side chain, and side chain-side chain interactions, respectively. The inter non-native interactions between A and B are given by

$$E^{NN}_{NB}(A,B) = \sum_{i=1}^{N_{NN}(A,B)} \epsilon_l\big(\frac{\sigma_i}{r_i}\big)^{6}, \tag{S8}$$

where $N_{NN}(A,B)$ is the total number of inter non-native interactions present between the beads of $A$ and $B$. The necessary parameters are listed in Table S1 and Table S2.

**Simulations:** We employed low friction Langevin dynamics simulations to effectively sample protein's conformational space and calculated important thermodynamic properties. The equation of motion for a bead with position coordinates $\vec{r_i}$ is depicted as,

$$m\ddot{\vec{r_i}} = -\zeta\dot{\vec{r_i}} + \vec{F_c} + \vec{\Gamma}, \tag{S9}$$

, where m is the mass of the respective protein bead, $\zeta$ is the friction co-efficient, $\vec{F_c} = -\frac{\partial E_{CG}(\{\mathbf{r}\},0,t)}{\partial \vec{r_i}}$, $\vec{\Gamma}$ is the random force with a white noise spectrum. The autocorrelation function of the random force in the discretised form is given by $\langle\Gamma(t)\ \Gamma(t+nh)\rangle = \frac{2\zeta k_B T}{h}\delta_{0,n}$, where $n = 0, 1, ...$ and $\delta_{0,n}$ is the Kronecker delta function. Velocity Verlet algorithm[10, 11] is used to integrate the equation of motion. We used $\zeta = 0.05\ m/\tau_L$ and time step, $h = 0.005\ \tau_L$, where $\tau_L$ is the unit of time used to advance the simulation.

To probe the folding kinetics of proteins, we used Brownian dynamics simulations where equations of motions are integrated using the Ermak-McCammon algorithm[12],

$$r_i(\vec{t+h}) = r_i(\vec{t}) + \frac{h}{\zeta}\vec{F_c} + \vec{\Gamma}. \tag{S10}$$

$\vec{\Gamma}$ is the random force with mean $\langle \Gamma(h) \rangle = 0$, and variance $\langle \Gamma(h)^2 \rangle = \frac{2k_B T h}{\zeta}$. The friction coefficient $\zeta = 31.9\ m/\tau_H$ is chosen close to that of water and integrating time step $h$ is set to $0.05\tau_H$. In the simulations, we chose the unit of length $a = 1$ Å, energy $\epsilon = 1$ kcal/mol, and mass $m = 1.8 * 10^{-22}$ g. Unit of time in Langevin dynamics simulations is $\tau_L = \sqrt{ma^2/\epsilon} = 0.51$ ps, while time step $\tau_H$ in Brownian dynamics simulations is $\tau_H \approx \frac{\zeta_H a^2}{k_B T} = \frac{(\zeta_H \tau_L/m)\epsilon}{k_B T}\tau_L \approx 53$ ps. We used weighted histogram analysis method (WHAM)[13] to compute average thermodynamic properties from Langevin dynamics simulations performed at a range of temperatures. Equilibrium properties like average internal energy $\langle E \rangle$, specific heat capacity $C_v (= \frac{\langle E^2 \rangle - \langle E \rangle^2}{k_B T^2}$, $k_B$ is the Boltzmann constant) and folding free energy $F$ are computed from the Langevin dynamics simulations. The detailed description of the WHAM equations to compute equilibrium properties is reported elsewhere[14]. Three-dimensional free energy $F(\chi_{Ntd}, \chi_{Ctd}, \chi_{Int})$ $\{= -k_B T \log P(\chi_{Ntd}, \chi_{Ctd}, \chi_{Int})\}$ is projected along fraction of native contacts present in Ntd ($\chi_{Ntd}$), in Ctd ($\chi_{Ctd}$) and in between Ntd and Ctd ($\chi_{Int}$) while two-dimensional free energy $F(\chi_{Ntd}, \chi_{Ctd})$ $\{= -k_B T \log P(\chi_{Ntd}, \chi_{Ctd})\}$ is projected along $\chi_{Ntd}$ and $\chi_{Ctd}$. Fraction of native contacts in any structural unit is defined as

$$\chi_{Str} = \frac{1}{N_N^{Str}} \sum_{i=1}^{N_N^{Str}} \left\{ \frac{1.0 - (\frac{r_i - r_{cry,i}}{r_0})^6}{1.0 - (\frac{r_i - r_{cry,i}}{r_0})^{12}} \right\} \tag{S11}$$

where $N_N^{Str}$ is the total number of native contacts present in the structural unit, $r_{cry,i}$ and $r_i$ are the distance between the $i^{th}$ pair of beads in the native structure and in a conformation given by $\{\mathbf{r}\}$, respectively, and $r_0$ is the cut-off distance. To track dimerization, number of native contacts $Q_{AB}$ are computed between monomer $A$ and monomer $B$, where contact between a pair of beads is computed as

$$Q = \left\{ \frac{1.0 - (\frac{r_i - r_{cry,i}}{r_0})^6}{1.0 - (\frac{r_i - r_{cry,i}}{r_0})^{12}} \right\}, \tag{S12}$$

which is similar to eq. S11. Symmetric Go potential (eq. S6) is used to define native contacts between monomer $A$ and monomer $B$.

Table S1: Energy function parameters for the SOP-SC model

| Parameters | Values |
|---|---|
| $R_o$ | 2.0 |
| $k$ | 20 kcal/(mol. Å$^2$) |
| $R_c$ | 8 Å |
| $r_0$ | 2 Å |
| $\epsilon_h^{bb}$ | 0.45 kcal/mol |
| $\epsilon_h^{bs}$ | 0.45 kcal/mol |
| $\epsilon_l$ | 1.0 kcal/mol |
| $\sigma^{bb}$ | 3.8 Å |

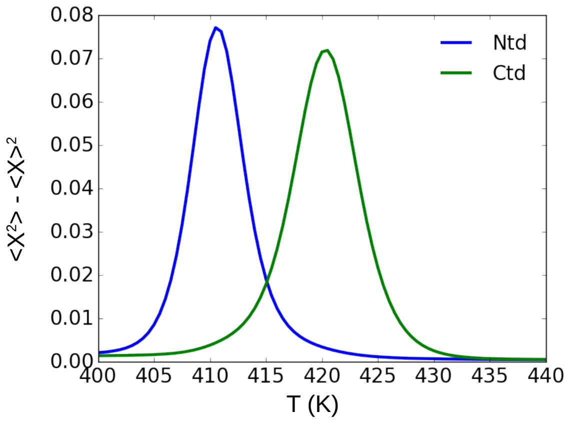


Figure S1: Deviation in fraction of native contacts ($\chi$) in Ntd and Ctd of H$\gamma$C Crys as a function of temperature ($T$). A two state folding is observed for both the truncated domains.

Table S2: SOP-SC model parameters for HγC Crys and HγD Crys

| Parameters | 2NBR | 1HK0 |
|---|---|---|
| $N_B$ | 345 | 345 |
| $N_N^{bb}$ | 577 | 565 |
| $N_N^{bs}$ | 1287 | 1280 |
| $N_N^{ss}$ | 554 | 541 |
| $N_N^{Ntd}$ | 1104 | 1093 |
| $N_N^{Ctd}$ | 1077 | 1080 |
| $N_N^{Int}$ | 161 | 143 |
| $N_N^{bb}(A,B)$ | 1154 | 1130 |
| $N_N^{bs}(A,B)$ | 2574 | 2560 |
| $N_N^{ss}(A,B)$ | 1108 | 1082 |
| $N_{NN}$ | 56235 | 56267 |
| $N_{NN}(A,B)$ | 114880 | 114944 |
| $N_{ang}^{bb}$ | 171 | 171 |
| $N_{ang}^{bs}$ | 344 | 344 |
| $N_c$ | 46 | 46 |

Table S3: Side-chain radii of amino acids

| Residue | Radius (Å) |
|---|---|
| Gly | 0.5 |
| Ala | 2.52 |
| Val | 2.93 |
| Leu | 3.09 |
| Ile | 3.09 |
| Met | 3.09 |
| Phe | 3.18 |
| Pro | 2.78 |
| Ser | 2.59 |
| Thr | 2.81 |
| Asn | 2.84 |
| Gln | 3.01 |
| Tyr | 3.23 |
| Trp | 3.39 |
| Asp | 2.79 |
| Glu | 2.96 |
| Hsd | 3.04 |
| Lys | 3.18 |
| Arg | 3.28 |
| Cys | 2.74 |

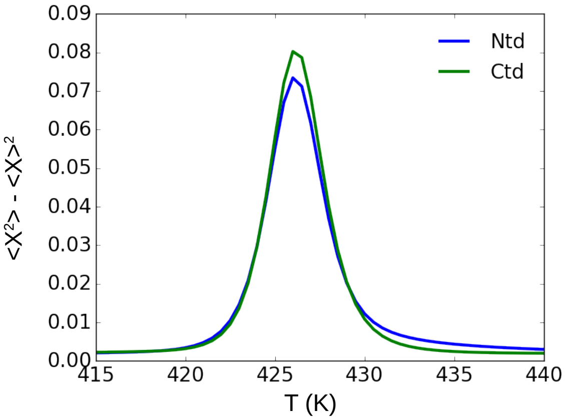


Figure S2: Deviation in fraction of native contacts ($\chi$) in Ntd and Ctd of H$\gamma$D Crys as a function of temperature ($T$). A two state folding is observed for both the truncated domains.

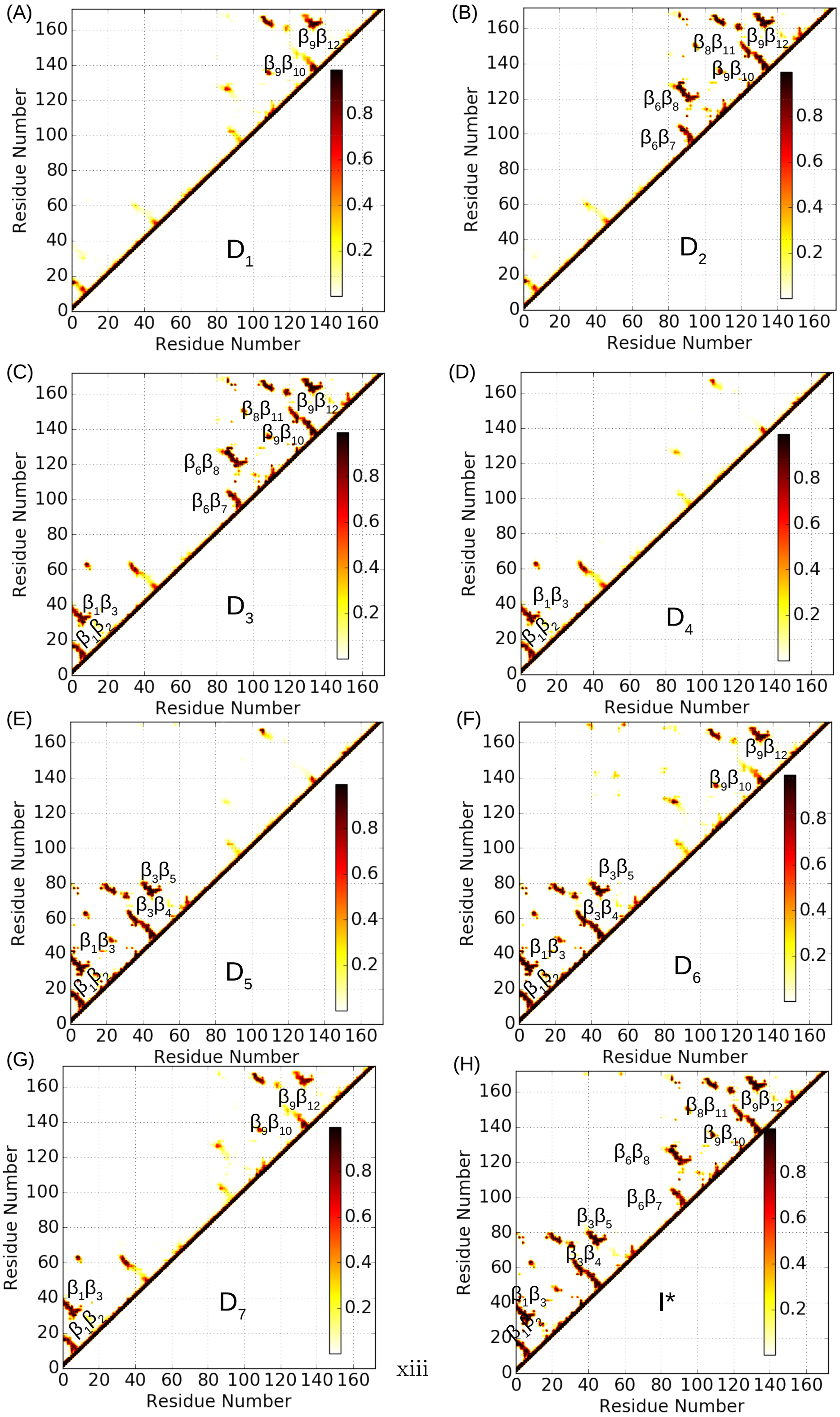




Figure S3: Contact maps of the intermediates ($D_1$, $D_2$, $D_3$, $D_4$, $D_5$, $D_6$, $D_7$ and $I^*$) populated in the folding FES of H$\gamma$D Crys. Contacts involving $\alpha$ helices and loops are not marked.

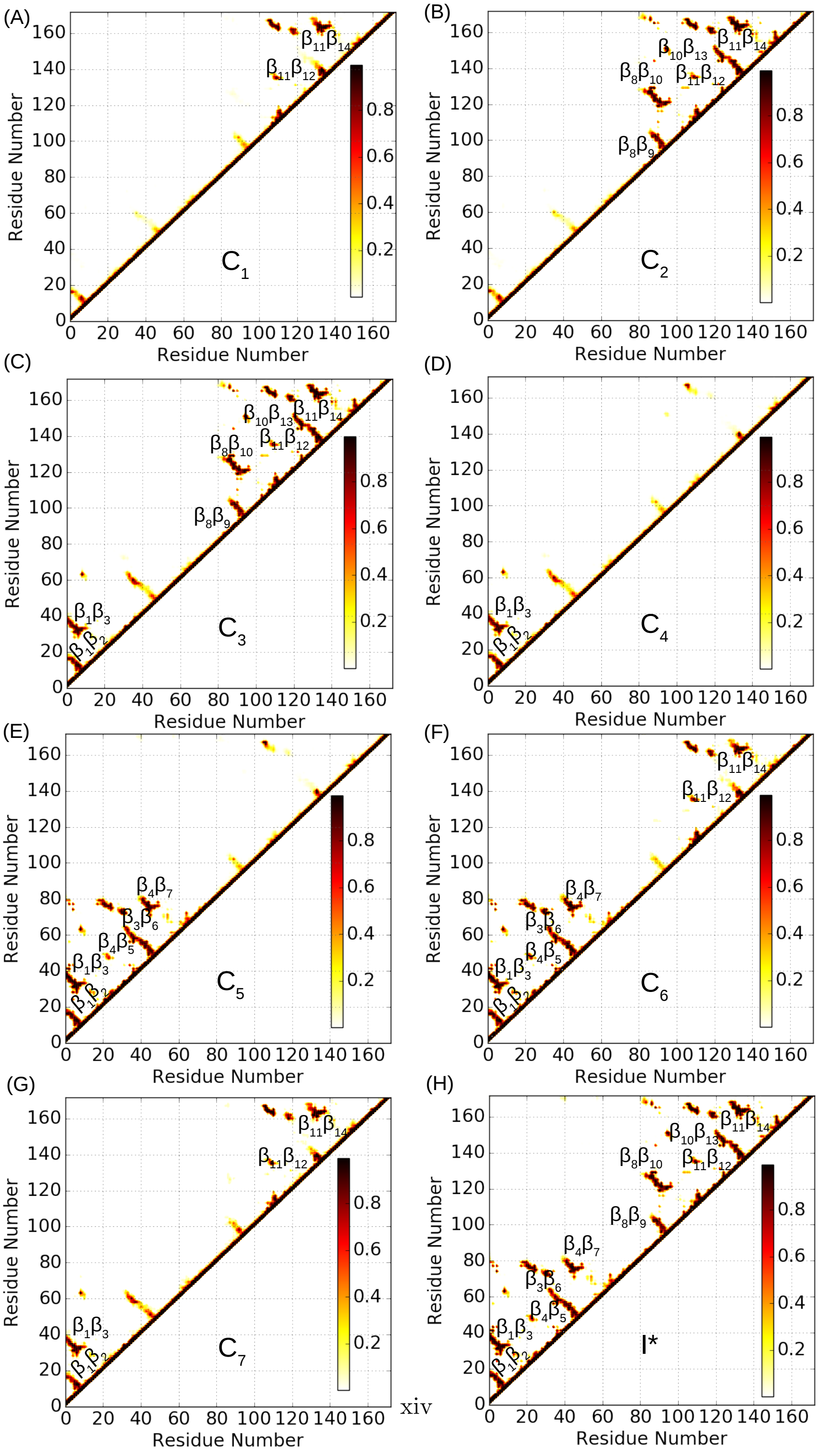




Figure S4: Contact maps of the intermediates ($C_1$, $C_2$, $C_3$, $C_4$, $C_5$, $C_6$, $C_7$ and I*) populated in the folding FES of H$\gamma$C Crys. Contacts involving $\alpha$ helices and loops are not marked.

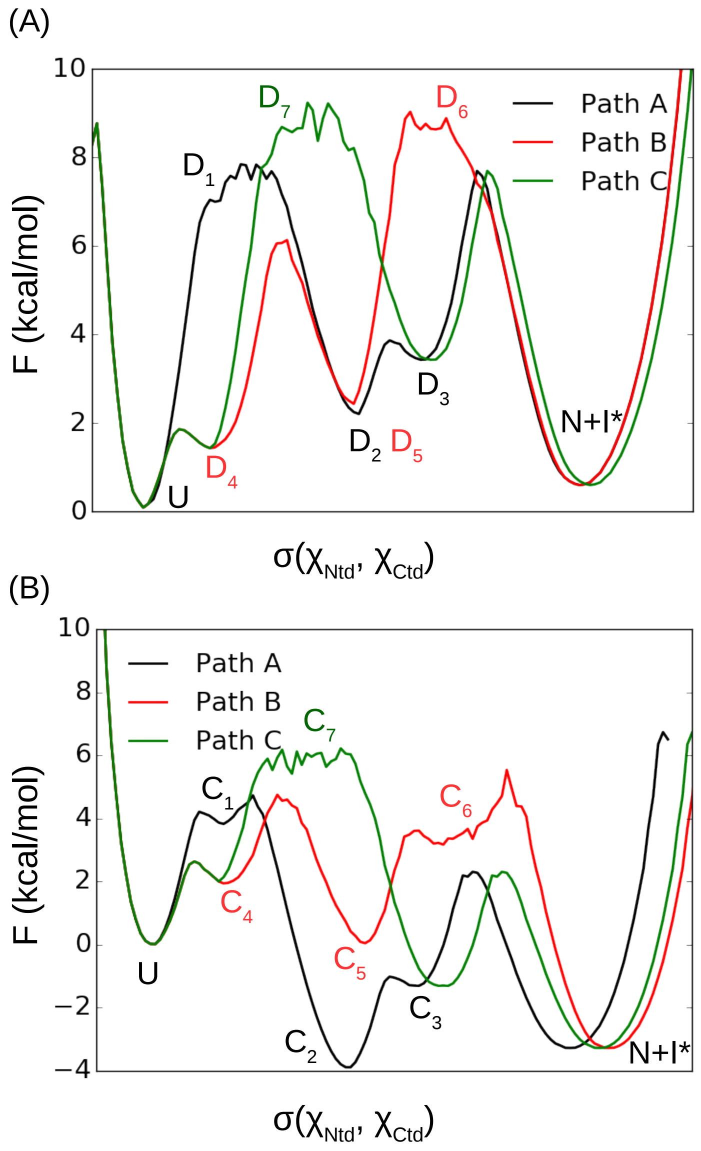


Figure S5: Minimum energy pathways $\sigma(\chi_{Ntd}, \chi_{Ctd})$ connecting the unfolded state $U$ to the folded state $N$ in (A) H$\gamma$D Crys and (B) H$\gamma$C Crys. $\sigma(\chi_{Ntd}, \chi_{Ctd})$ is computed from the two-dimensional FES $F(\chi_{Ntd}, \chi_{Ctd})$ (Figure 1E,2D) using the software MEPSA[15].

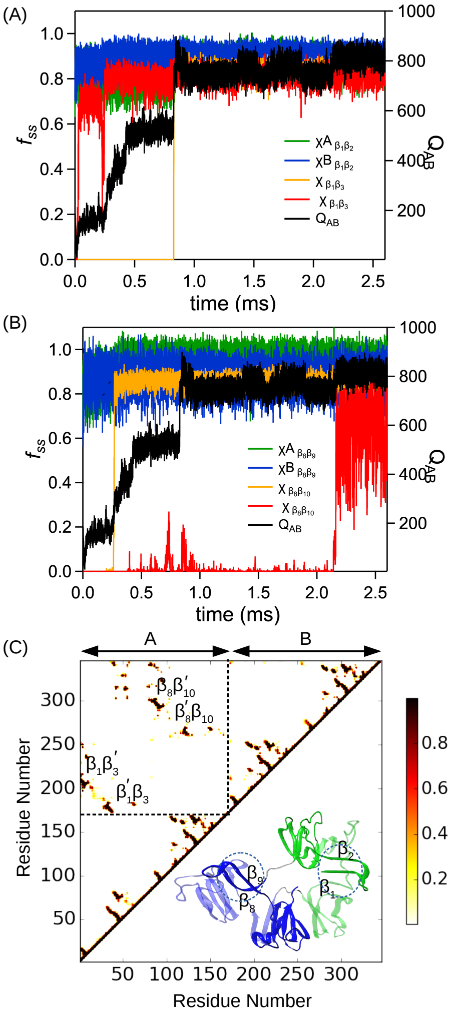


Figure S6: Multistep dimerization in H$\gamma$C Crys while refolding at $T$ = 300 K. Secondary structural elements from Ntd ($\beta_1\beta_2$) and Ctd ($\beta_8\beta_9$) get swapped leading to a double domain swapped structure. (A) Time evolution of the fraction of secondary structural contacts between $\beta_1\beta_2$ strands in monomer A, B ($\chi^A_{\beta_1\beta_2}$, $\chi^B_{\beta_1\beta_2}$ ) and between $\beta_1$ strand of monomer A and $\beta'_3$ strand of monomer B ($\chi_{\beta_1\beta'_3}$) and vice versa ($\chi_{\beta_1\beta'_3}$) along with number of native contacts formed between monomer A and B ($Q_{AB}$). (B) Time evolution of fraction of secondary structural contacts between $\beta_8\beta_9$ strands in monomer A and B ($\chi^A_{\beta_8\beta_9}$, $\chi^B_{\beta_8\beta_9}$ ) and between $\beta_8$ strand of monomer A and $\beta'_{10}$ strand of monomer B ($\chi_{\beta_8\beta'_{10}}$) and vice versa ($\chi_{\beta_8\beta'_{10}}$). (C) Contact maps of the final domain swapped dimer structures. $\beta_1$ strand makes contacts with $\beta'_3$ strand of the other monomer and vice versa after swapping of $\beta_1\beta_2$ strands. The first hairpin loop of $M_3$ ($\beta_8\beta_9$) get swapped between the Ctds while $\beta_8$ strand forms contact with $\beta'_{10}$ strand of the other monomer and vice versa.

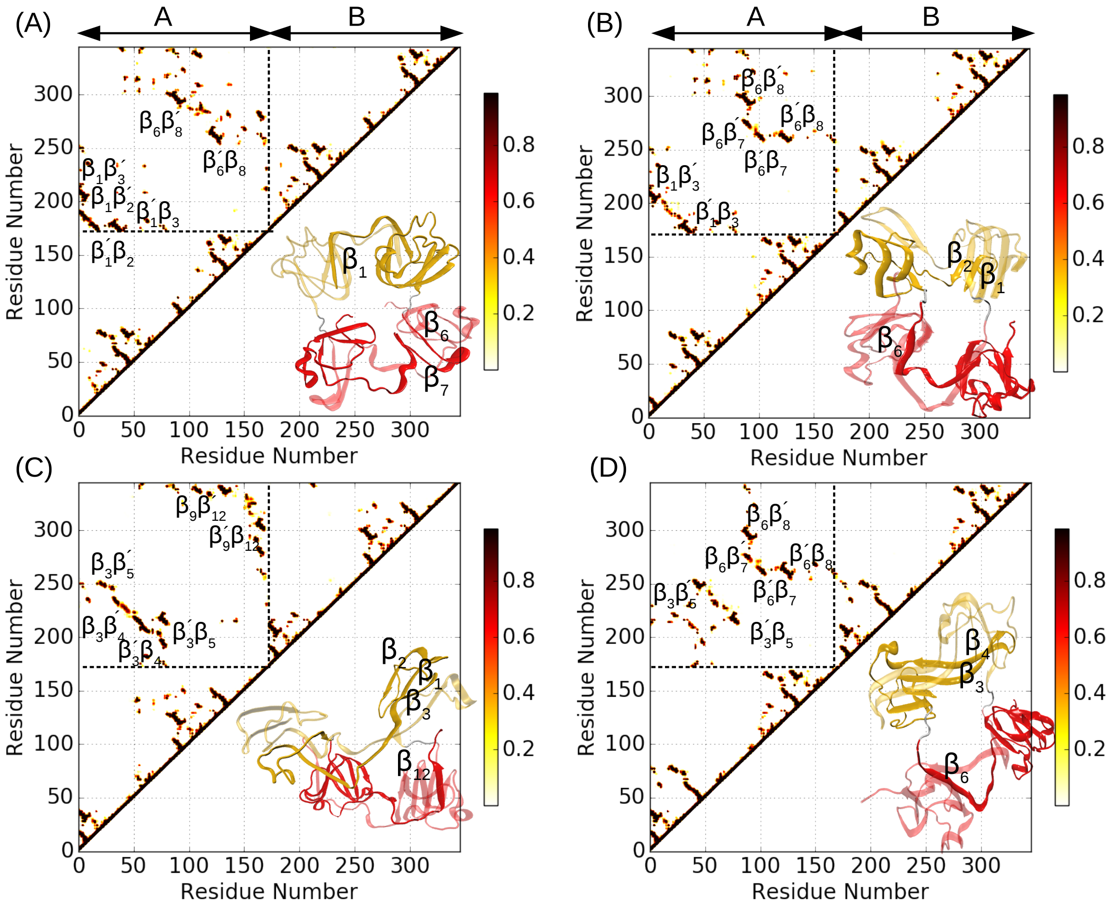

Figure S7: Contact map and representative structures of dimers (population $< 5$ %) obtained during refolding of H$\gamma$D Crys.

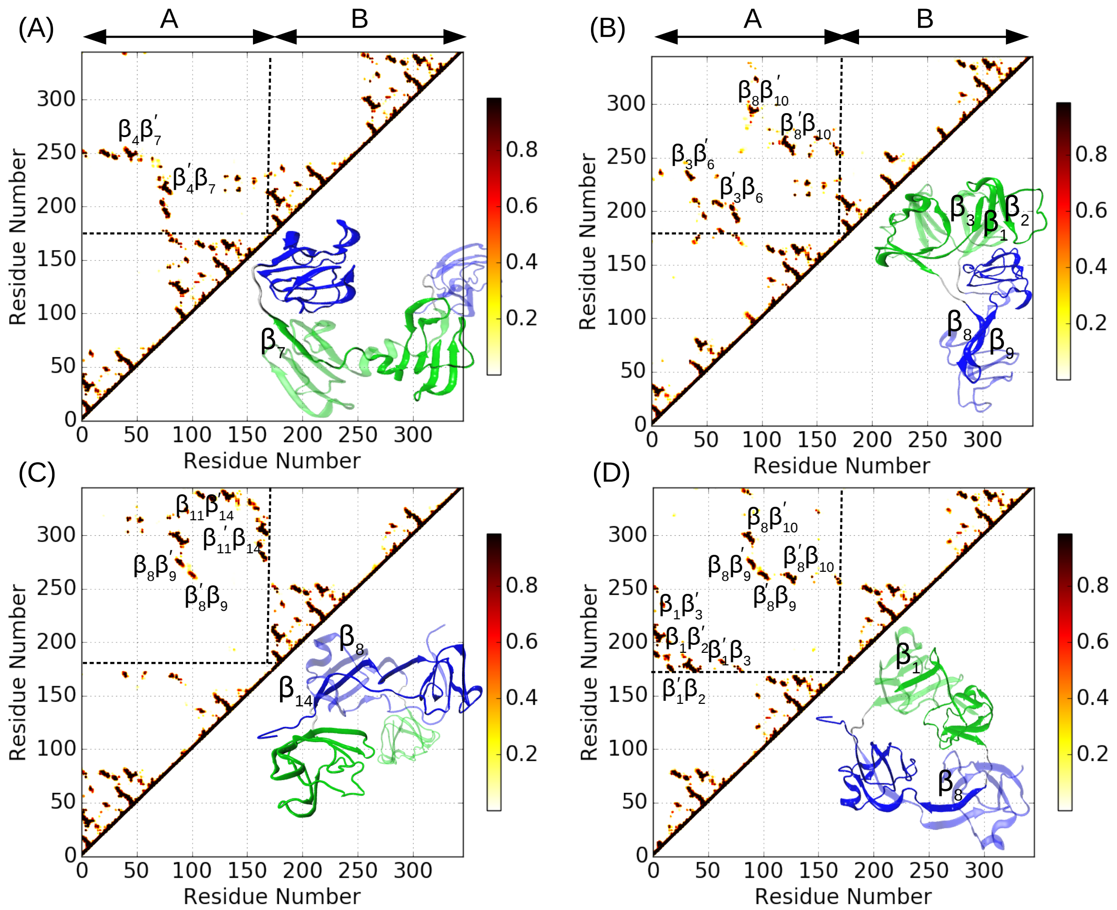


Figure S8: Contact map and representative structures of dimers (population < 5 %) obtained during refolding of H$\gamma$C Crys.

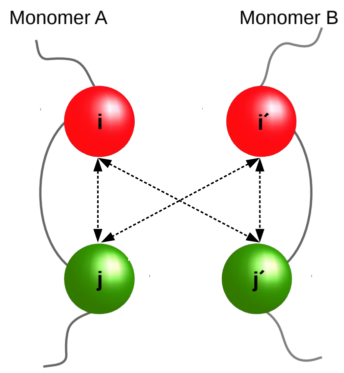


Figure S9: Symmetric Go potential acting between monomer $A$ and $B$. Native interactions present between residue $i$ and $j$ of monomer $A$, will also manifest into native interactions between residue $i'$ and $j'$ of monomer $B$, between residue $i$ and $j'$, and between residue $i'$ and $j$.
